# Transcriptional changes in non-human primate tissues after intrathecal delivery of serotype 9 adeno-associated viral vector: insights into organ toxicities

**DOI:** 10.1101/2025.05.09.652903

**Authors:** Fumiaki Aihara, Matthew Mardo, Vera Ruda, Tony Del Rio, Karine Bigot, Jochen Singer, Megumi Onishi-Seebacher, Philippe Couttet, Keith Mansfield, Eloise Hudry

## Abstract

Adeno associated virus (AAV)-based gene transfer has brought transformative therapeutic benefits to patients with otherwise untreatable genetic diseases. However, treatment-related organ toxicities, particularly for high doses, remain a safety concern in the clinic with some translatability in preclinical species. In the present study, we conducted an RNAseq analysis in non-human primates administered intrathecally with scAAV9-CBA-GFP, empty viral capsid particles, or a ‘Promoterless’ vector. This analysis revealed a broad and long-lasting (4 weeks after dosing) transcriptional impact of the viral transduction/transgene expression on tissues. Liver and DRG, known to be primary sites of toxicity induced by AAV9, had the highest viral load and the most significant transcriptional changes. Our analysis revealed that most of the differentially expressed genes were upregulated, and common gene signatures belonged to immune pathways (innate and adaptive), demonstrating a persistent low-grade immune response up to four weeks post-dosing. Interestingly and across all tissues considered, the impact of empty capsids or of the Promoterless vector was minimal, suggesting that both the presence of the capsid and of a productive viral genome contributed to the observed changes. This study provides a unique insight into the transcriptional responses to AAV9 in key tissues primarily exposed by the vector.

## INTRODUCTION

During the initial groundwork conducted in the 1980’s, Adeno Associated Viruses (AAV) have been speculated as potential vectors for introducing functional genes to target cells^1,2^. Due to their relative innocuity to the human population, low immunogenicity and broad tropism for mammalian cells, AAV viral vectors have become a vessel of choice for the transfer of genetic material. After decades of virological research, optimization in vector genetic payload, and manufacturing^3,4^, the field has finally progressed to the point where AAV-based gene therapies have gained approval for the treatment of several monogenic disorders, with seven approved products in 2024^5^. A meta-analysis of AAV-based gene therapies clinical trials show a high number of trials meeting primary efficacy endpoints and safety requirements, in particular the most common intravenous administration (80% of 48 trials) as well as intrathecal (100% of 4 trials)^6^.

Despite those initial successes, several challenges still hinder the broad expansion of this therapeutic modality. In particular, the emergence of organ toxicities in patients (hepatotoxicity, cardiotoxicity and potentially dorsal root ganglia toxicity after intravenous or intrathecal administration, RPE toxicity after subretinal dosing, etc.) has generally impacted the benefit/risk balance for AAV gene therapies^7,8^. In the context of systemic administration for the treatment of non-hepatic disorders, therapeutic benefit can often only be achieved after delivering high doses of viral vector, therefore increasing the risk for bystander tissue toxicity. As such, rises in biomarkers of organ toxicity have been observed in patients^7,9–12^, resulting in asymptomatic subacute injury or in severe adverse events and even death^13–16^. In ongoing clinical trials in 2022, 16 trials with IV dosing and 11 trials with IT dosing have reported test article related adverse effects^6^.

As a non-clinical correlation, previous work published by our group and others have aimed at investigating AAV-induced organ toxicities in non-human primates (NHPs)^10,17–23^. Our retrospective analysis across several primate studies revealed that the biodistribution of scAAV9 vector genomes is generally highest in the liver and dorsal root ganglia, two tissues in which safety findings are generally reported. In liver, we have observed sharp dose response for AAV-driven toxicity, with low (IT dosing up to 1.00×10^13^ vg/NHP) to moderate (IV dosing at ∼1×10^14^vg/kg) systemic exposure presenting asymptomatic elevation of liver enzymes, while higher doses administered intravenously (∼2×10^14^vg/kg) led to clinical signs associated with severe adverse hepatotoxicity. Interestingly, NHPs that received scAAV9 empty capsids or Promoterless genome intrathecally showed no elevations in transaminase levels or histological changes in liver, indicating capsid related immune responses cannot solely trigger AAV-induced hepatotoxicity. Molecular localization of select NHPs afflicted with AAV-induced hepatotoxicity revealed an increase of macrophage markers, and proteins associated with the RIG-I pathway. Such findings were generally aligned with the work recently published by Hordeaux *et al.*^12^ which reported acute thrombocytopenia after administration of high dose AAV in NHPs, as well as the presence of anti-AAV antibodies 7 days post dose. Of the animals that developed acute thrombocytopenia, the majority had recovered, while a few progressed to liver failure. When examining peripheral neuropathy driven by AAV viral vectors, we also described the severity, heterogeneity and timeline of histopathological changes observed in the primates’ DRGs. Such toxicity generally correlated with circulating neurofilament light chain (NfL) in the cerebral spinal fluid (CSF) and blood, highlighting the usefulness of such biomarker to follow AAV-driven peripheral neuropathy in this nonclinical species^11^. Altogether, these published studies highlight the usefulness of nonclinical assessment to further understand AAV-driven organ toxicities, providing important insight into the timeline, clinical variability and underlying pathophysiology of such effects.

In this context, the nonclinical investigation and mitigation of such toxicities will be a key step to further the development of AAV gene therapies. To gain insight into the underlying molecular mechanisms, we conducted a retrospective transcriptomic analysis focused on two non-human primate studies using a tool AAV9 vector expressing the reporter transgene Green Fluorescent Protein delivered into the intrathecal space^10^. Our investigation revealed a clear distinction in transcriptional responses in the DRG and liver between fully functional and non-functional AAV (empty capsids or Promoterless). Most upregulated genes were immune-related and combined innate- and adaptive immune changes. Interestingly, while the ‘Promoterless’ viral vector did not cause any evidence of organ toxicity from a pathological standpoint, transcriptional responses following the administration of this test article were closer to the fully functional AAV than the empty viral capsids.

## RESULTS

### In-life summary and organ toxicities (primarily liver and DRGs) observed after intrathecal dosing with AAV9 in the Cynomolgus macaques

Intrathecal dosing with AAV9 full particles (scAAV9-CBA-GFP), empty AAV9 capsids or Promoterless AAV9 at doses of 1 ×10^13^ to 3×1013vg/NHP was generally tolerated with minimal clinical signs observed. In study A, NHPs administered high dose scAAV9-CB-GFP showed elevated levels of bilirubin and liver enzymes that correlated with transient symptoms such as whole-body tremor, abnormal yellow periorbital color, and general decreased activity. In study B, elevated levels of alanine aminotransferase (ALT) were also observed around Day 15 post-dosing, without any association with clinical signs (Table S1). At necropsy (Day 28), microscopic findings observed in the hepatic tissue consisted in single cell necrosis, oval cell hyperplasia and mononuclear cell infiltration, therefore demonstrating. Single-cell necrosis and neuroinflammatory changes were also observed in DRGs, which correlated with elevated levels of NfL that peaked on Day 28. All hepatic and DRG toxicities were only detected after administration of full AAV9 viral particles, but not empty capsids or Promoterless test articles^8,14^. No remarkable changes were detected in heart, skeletal muscle and spleen, the other tissues included in our transcriptomics analysis. The vector biodistribution of the viral genome was also highest in liver and DRGs as compared to the other organs evaluated (Figure S1A)^10^, therefore suggesting that such test-article related changes were mostly driven by exposure of the tissues to the full viral vector. By contrast to the biodistribution, the transgene expression level was not highest in liver and DRGs (Figure S1B, C) and higher counts per million (CPM) of the transgene were detected in the heart and skeletal muscle from both study A and B (Figure S1B, C). This discrepancy between vector genome biodistribution and transgene transcript level suggested that while both the capsid and an ‘active’ payload are necessary to compromise the tissues’ integrity (liver and DRGs), the amount of transgene product itself was not directly predictive of the observed toxicities (even when expressing a foreign protein such as GFP).

### Lasting transcriptional changes observed across key tissues transduced by self-complementary AAV9 4 weeks after intrathecal dosing in Cynomolgus macaques

An initial evaluation of AAV9’s impact on a subset of tissues (liver, DRG, heart, skeletal muscle and spleen) was performed by comparing the number of differentially expressed genes across tissues from each treatment group (full scAAV9-CBA-GFP particles, empty viral particles and Promoterless vector) with the concomitant vehicle-treated group. In general, and in alignment with the anatomic pathology changes observed, the extent of the transcriptional changes (number of genes significantly up and downregulated) was more pronounced in both DRG and in the liver from animals dosed with the AAV9-full vector compared to the group given empty viral particles. Plotting the distribution of all upregulated genes across all treatment groups showed a clear increase in the number of genes above 1.0 log_2_FC in NHPs that have received fully functional AAV in DRG, and liver (Figure 1).

**Figure 1.**
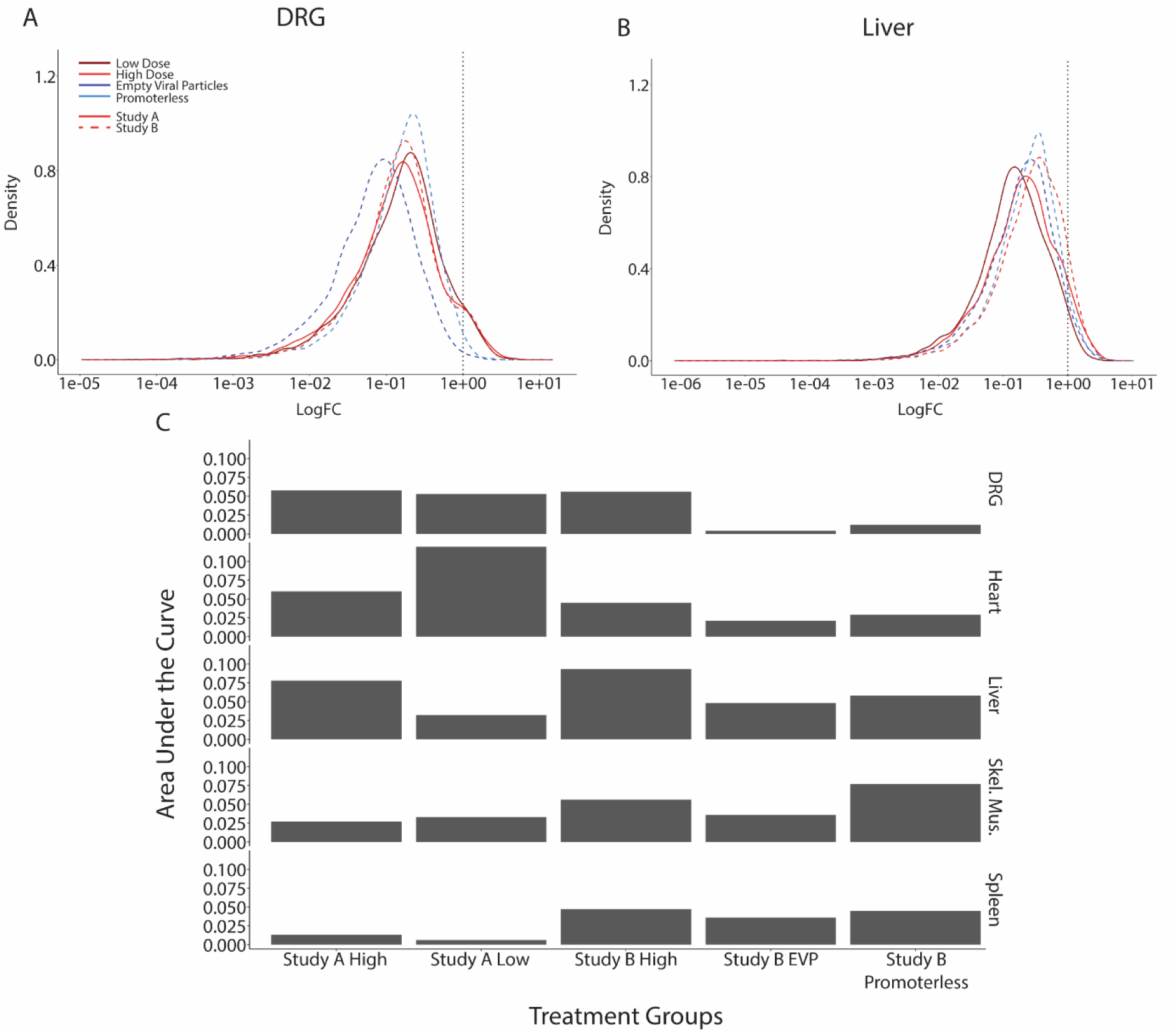
Density plots from upregulated genes in DRG and liver after IT administration of fully functional scAAV9-CBA-GFP vector, empty capsids or Promoterless vector. Density plots of the distribution of all upregulated genes from A) DRG or B) liver. The black dotted vertical line indicated Log_2_FC > 1.0. The red shaded lines represent data from animals dosed with the fully functional AAV. The solid or dotted curves indicate groups from study A or study B, respectively. C) Bar graph representing the area under the curve of log_2_FC >1.0 for each treatment arm and organ.

Across all the different tissues analyzed, more significantly upregulated genes were detected in DRGs and the amplitude of such changes was greater from animals dosed with the fully functional AAV particles compared to empty capsids or Promoterless vector (Figure 2). The number of downregulated genes was, by contrast, lesser. In the experimental groups that received the fully functional AAV across both studies, over 300 genes were found to be significantly upregulated while only up to 50 genes were downregulated. Those results reflect broad transcriptional changes in DRGs at the time of necropsy (4 weeks post-dosing), which generally coincides with the peak of NfL increases in CSF and blood and with the observed histopathological changes (more severe between 3-6 weeks) after intrathecal dosing^11^.

**Figure 2.**
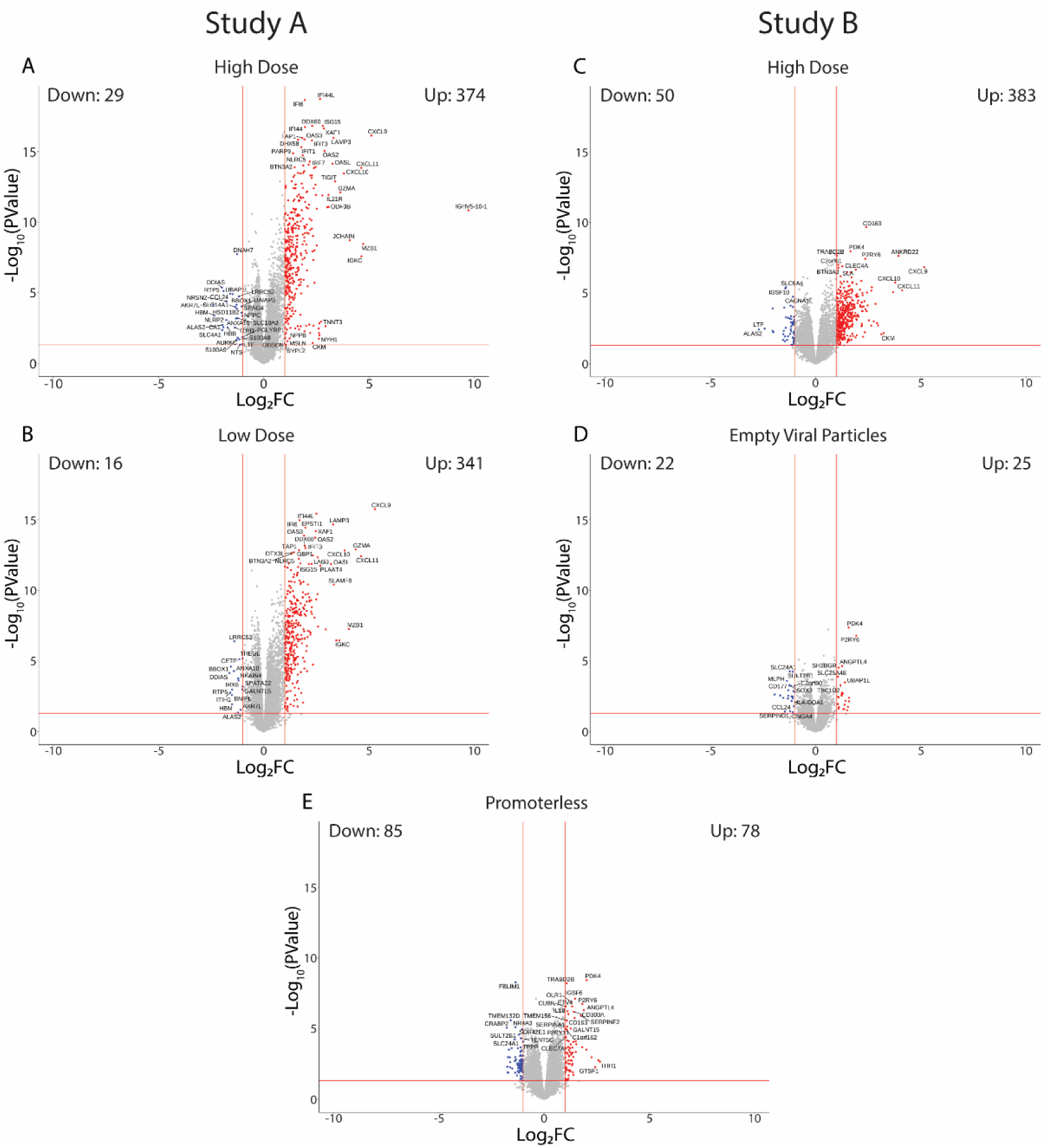
Volcano plots representing up- and down-regulated genes in DRGs after IT delivery of scAAV9-CBA-GFP, AAV9 empty capsid and Promoterless vector. A) and B) represent the DEGs identified in study A after administration of a low (A) or high (B) dose of fully functional AAV9. The volcano plots on the right panel represent the up- and down-regulated genes in DRGs exposed to scAAV9-CBA-GFP (fully functional, C), empty viral particles (D) or the Promoterless vector (E). Vertical red lines indicate the threshold of Log_2_FC of 1. The horizontal red lines indicate Log_10_(P value) of 0.05.

In liver, a relatively equivalent number of genes were found to be upregulated and downregulated upon scAAV9-CBA-GFP IT dosing as compared with hepatic tissue from the control groups (Figure 3A-C). In Study A, 173 genes were upregulated, and 105 genes were significantly downregulated for animals dosed at 1×10^13^vg/NHP. Those numbers sensibly increased in the high dose group (3×10^13^vg/NHP) from the same study (447 genes upregulated and 125 genes down-regulated) (Figure 3B), therefore suggesting that the amplitude of the transcriptional change is dose-responsive (when comparing experimental groups within the same study). For study B, 504 were upregulated while 539 genes were down-regulated in the liver from animals administered with the full scAAV9CB-GFP as compared to vehicle control (Figure 3C). Our previous report showed peak elevation of transaminases in plasma at day 15 followed by a drop at day 28 (day of termination) after intrathecal dosing with self-complementary AAV9 (at a later timepoint than the changes generally observed post-systemic dosing, which tend to peak within the first week post-dose), and therefore the transcriptional changes observed at 4 weeks may be relatively posterior to the initial liver injury^10^. In study A, we observed the low dose groups have shown ALT levels equal to or higher than the high dose groups. Our transcriptional analysis appears to reflect this with more significantly upregulated genes in the low dose group^10^. The liver volcano plots generated from NHPs that have received Promoterless or empty capsids AAV particles presented with lesser overall changes as compared to the impact observed with the full viral particles within the same study (similarly as with the DRGs). In the empty viral capsids, 279 upregulated genes and 180 down-regulated genes were found to be significantly altered in comparison to vehicle control (Figure 3D). The Promoterless group showed more global transcriptional changes compared to the empty viral capsid group with 286 upregulated and 573 genes downregulated (Figure 3E). The observed impact of the empty capsids and Promoterless vector on the liver transcriptional profile suggests that each element of those control test articles (capsid only, capsid plus an inactive payload) is inducing long-lasting changes on the hepatic tissue. That said, no elevations of transaminases were detected in those experimental groups, therefore suggesting that such transcriptional changes may not be clinically impactful.

**Figure 3.**
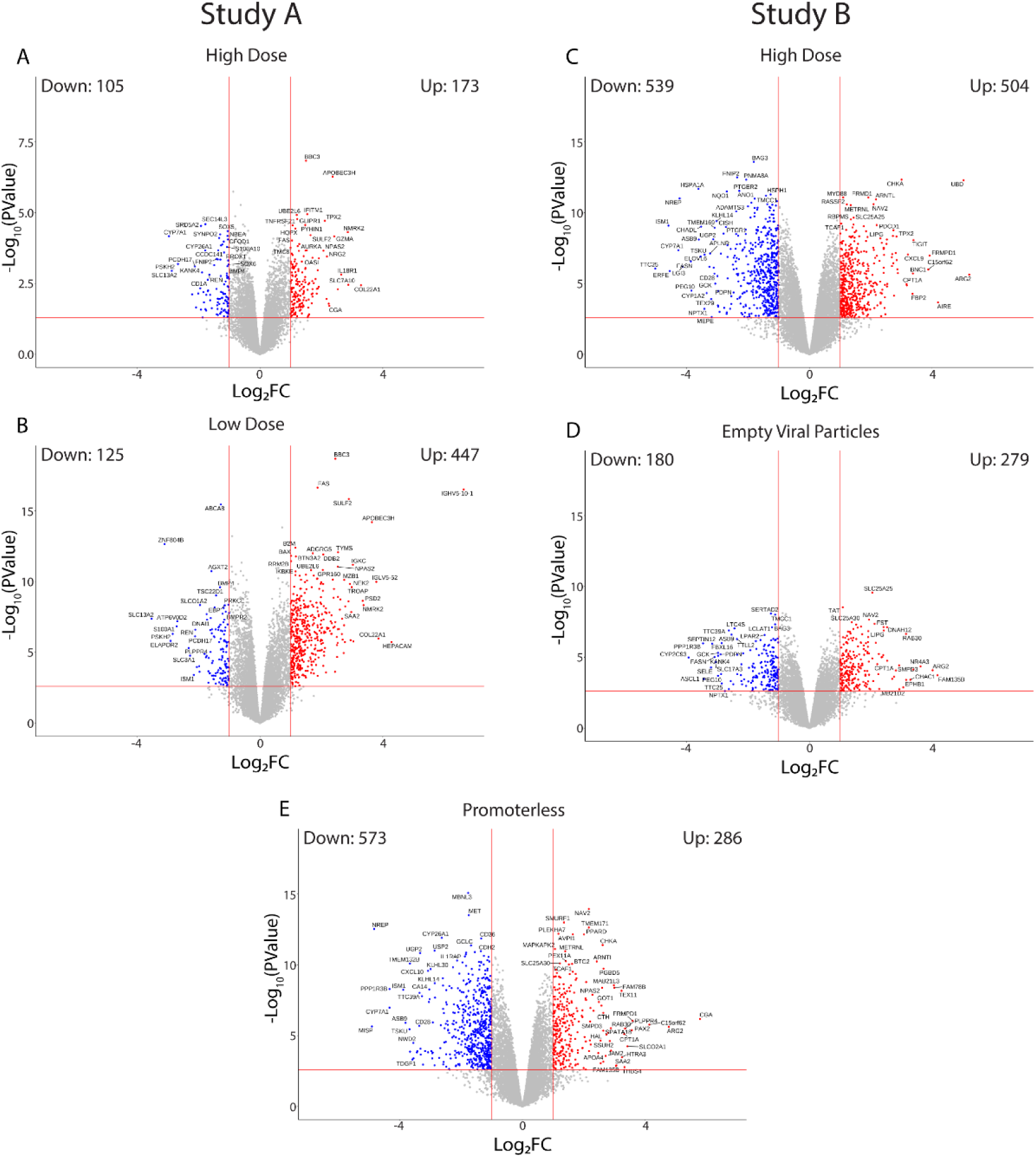
Volcano plots representing up- and down-regulated genes in liver after IT delivery of scAAV9-CBA-GFP, AAV9 empty capsid and Promoterless vector. A) and B) represent the DEGs identified in study A after administration of a low (A) or high (B) dose of fully functional AAV9. The volcano plots on the right panel represent the up- and down-regulated genes in DRGs exposed to scAAV9-CBA-GFP (fully functional, C), empty viral particles (D) or the Promoterless vector (E). Vertical red lines indicate the threshold of Log_2_FC of 1. The horizontal red lines indicate Log_10_(P value) of 0.05.

Another tissue of interest selected for our analysis was heart, considering AAV9 distributes to the cardiac tissue and that AAV-driven cardiotoxicity has been reported in clinical and pre-clinical studies^24–26^. Symptoms of myocarditis have been observed in gene therapy trials for Duchenne muscular dystrophy, including elevated troponin I levels, focal wall-motion abnormalities, and decline of the left ventricular ejection fraction across several trials^15,24,26^. In the heart, the observed biodistribution of AAV9 (vector genome per diploid genome) is generally lower levels than other peripheral tissues such as liver or DRGs^10^, but interestingly the transgene expression levels and amounts of transgene protein in the cardiac tissue are very elevated (by contrast to the liver, which shows high viral copy numbers but relatively low transgene product levels)^27^. In study A, clinical pathology changes revealed increased levels of CK in study A on day 15 in the high dose group (mean FC: 6.6), which correlated with microscopic changes in that tissue consisting of minimal to moderate mononuclear cell inflammation (degeneration of cardiomyocytes surrounded by mononuclear cells). Signs of cardiac toxicity were more subdued in study B with increased CK levels observed in only two animals dosed with the full scAAV9-CBA-GFP vector on Day 28 while such an increase was observed in one NHP dosed with empty particles and one NHP dosed with vehicle control. In that study (B), the anatomic pathology evaluation of the heart tissue was also found to be unremarkable. Interestingly, a larger number of differentially expressed genes were detected in study A compared to study B for animals dosed with the fully functional AAV9 (comparing Figure 4A-C), potentially reflecting the stronger effect of the viral vector in study A (for which clinical pathology and histology changes were observed). In the high dose group from study A, there were 111 down-regulated genes and 305 upregulated genes (Figure 4A). For the low dose group, we observed 187 downregulated and 642 upregulated genes compared to the vehicle control (Figure 4B). By contrast, only 135 downregulated and 196 upregulated genes were detected in study B for animals dosed with the fully functional AAV (Figure 4C). A smaller impact was observed in animals that received empty viral particles or the Promoterless vector, with 70 or 114 downregulated genes and 76 or 121 upregulated genes detected in those two groups, respectively (Figure 4D, E). Such differences once again suggest that the long-lasting transcript transcriptional effect of AAV is associated with the capsid and a functional genome.

**Figure 4.**
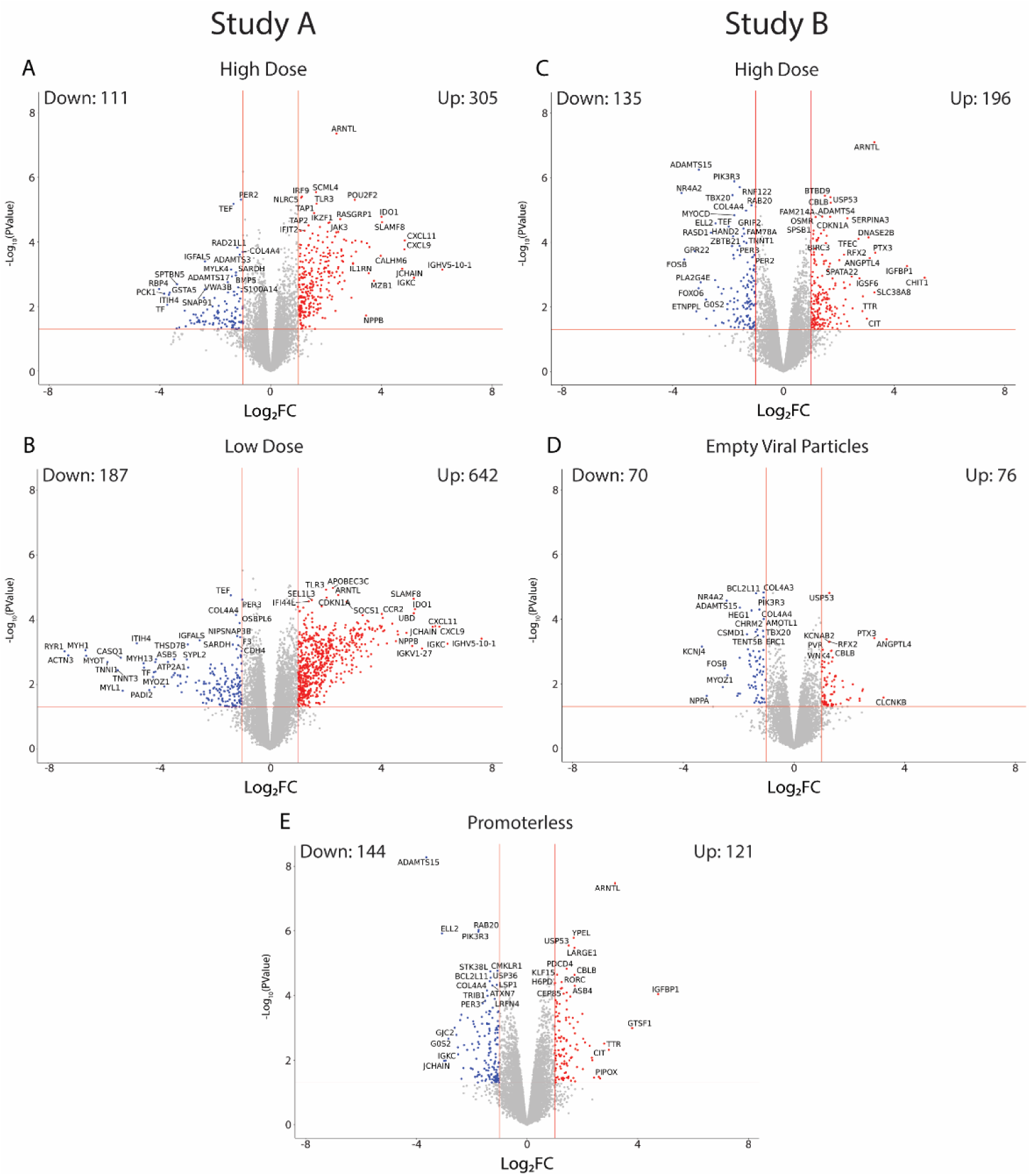
Volcano plots representing up- and down-regulated genes in heart after IT delivery of scAAV9-CBA-GFP, AAV9 empty capsid and Promoterless vector. A) and B) represent the DEGs identified in study A after administration of a low (A) or high (B) dose of fully functional AAV9. The volcano plots on the right panel represent the up- and down-regulated genes in DRGs exposed to scAAV9-CBA-GFP (fully functional, C), empty viral particles (D) or the Promoterless vector (E). Vertical red lines indicate the threshold of Log_2_FC of 1. The horizontal red lines indicate Log_10_(P value) of 0.05.

The transcriptional changes observed in skeletal muscle were generally fewer and of smaller magnitude than the observed changes in other tissues described above (Figure S2). Finally, and across all tissues considered, the spleen presented with the fewest transcriptional changes after intrathecal dosing with scAAV9 (Figure S3B). The smaller impact observed across both tissues (skeletal muscle and spleen) is generally aligned with the absence of remarkable findings detected from those organs at necropsy, even though this provides evidence that AAV transduction and transgene expression could cause some transcriptional effects in organs that do not show any signs of toxicity.

### Pathway analysis of AAV9-dependent transcriptional changes reveals common immunological responses four weeks after intrathecal delivery in the primate

Genetic expressions and biological functions are interwoven in a complex web of upregulation, downregulation, degradation, and feedback loops. To get some initial insight into the biological functions primarily impacted by AAV transduction, we conducted a pathway analysis on the sets of Differentially Expressed (DE) genes between AAV- and vehicle-dosed animals across both studies A and B for DRG and liver. As advised by the developers of the edgeR package, we used a p-value threshold of less than 0.0001 for significance^28^. The results of each intra-study pathway analysis (treatment groups versus control for each study) were then compared across studies to identify common pathways (Table S2-6). Through our analysis, 49 pathways were significantly upregulated in the DRG in both high-dose groups (p-values below 0.0001), while only 2 positively regulated pathways identified liver for the same groups. No downregulated pathways across all tissues were identified below this threshold. We filtered for common pathways that were significant in the high dose group (full AAV only) in both studies.

In the DRG, the top five most significant pathways associated with the high-dose group of AAV9 were “Influenza A”, “Epstein-Barr virus infection”, “Th17 cell differentiation”, “Lipid and atherosclerosis”, and “NOD-like receptor signaling pathway” (Table S2). In the “Influenza A” pathway, the most significantly expressed gene was *CXCL10* (Figure 5A). Some genes in the same pathway were related to viral RNA binding such as *OAS2*, *OAS3*, *DDX58*, and *TLR7*. Others were related to signal transduction and regulation, such as *IKBKE*, *IRF7*, *IRF9*, and *RSAD2*. Similar classes of genes also appear in the “Epstein-Barr virus infection” pathway, in addition to other genes of interest such as HLA genes (indicative of active antigen presentation) (Figure 5B). *CD3D*, *CD3G,* and *CD3E,* were identified across several pathways (“Th17 cell differentiation”, “Epstein-Barr virus infection), hinting at the increased presence of T-cells in the DRGs after AAV9 dosing (Figure 5B, C). In “Lipid and atherosclerosis” pathway, the most highly expressed genes were also largely related to immunological functions (*OLR1*, *FASLG* associated with apoptosis, *POU2F2* encoding a protein that binds to immunoglobulin gene promoters, and *VAV1*, an important gene for the development T-cell and B-cells) (Figure 5D). Under “NOD-like receptor signaling pathway” the cluster with the highest log_2_FC values contained genes we have observed in previously mentioned pathways (*OAS* family, *CASP* family, *IRF7*). The more unique genes in this pathway were *GBP1, 2,* and *3*, which are known to be induced by interferon signaling (Figure 5E).

**Figure 5.**
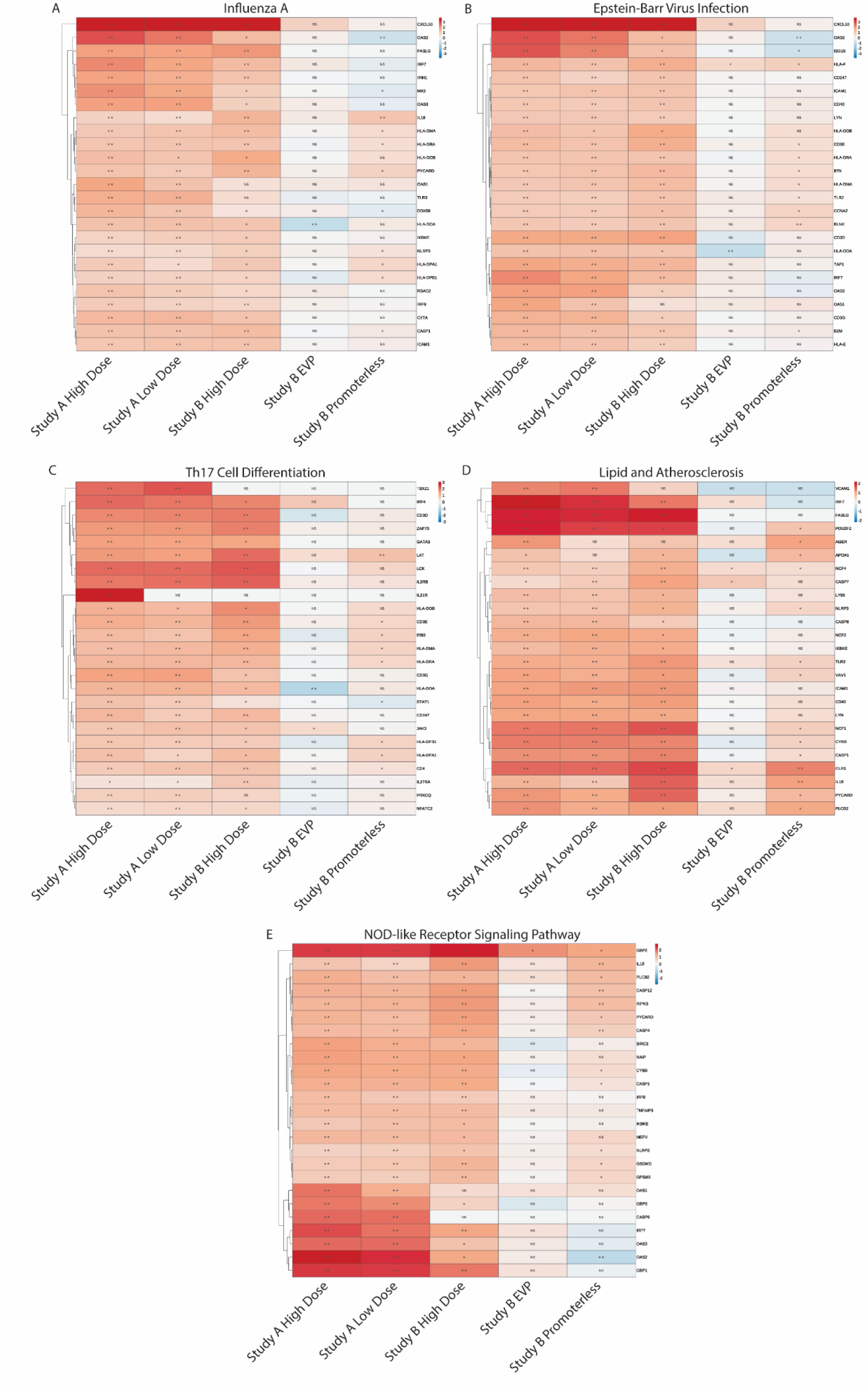
Top 5 most relevant pathways identified from differentially expressed genes identified in DRG. The 25 most significant genes associated for each pathway are being displayed on the right of each table, segregated by study arm. The most relevant pathways up-regulated with the administration of scAAV9-CBA-GFP (fully functional vector) IT were: ‘Influenza’ (A), ‘Epstein-Barr Virus Infection’ (B), ‘Th17 Cell Differentiation’ (C), ‘Lipid and Atherosclerosis’ (D) and ‘NOD-like Receptor Signaling Pathway’ (E). A few genes across both pathways were found to be also associated with the administration of the Promoterless vector, but not with the administration of empty capsids. The heatmap colors represent log_2_FC and the dendrograms on the right depict hierarchical clustering. NA: Not Applicable; NS: Not Significant; *: p<0.05; **: p<0.01; ***: p<0.001.

In the liver, fewer pathways were identified, and none significantly differed between animals dosed with empty viral capsids or vehicle control. Two pathways showed significance with a p-value below the 0.0001 when comparing the fully functional scAAV9 and the vehicle control. Those pathways were identified as “Antigen processing and presentation” and “Human T-cell leukemia virus 1 infection” (Table S4). Under the “Antigen processing and presentation” category, many of the upregulated genes belonged to the HLA genetic family, *TAP1*, *TAP2* and *CD27* (Figure 6A). Within the “Human T-cell leukemia virus 1 infection” pathway, many genes clustered with the group dosed with functional AAV are genes seen in previous pathways (*HLA* genes, *CD3*, *CD4*, *JAK3*, and *IL15)* (Figure 6B).

**Figure 6.**
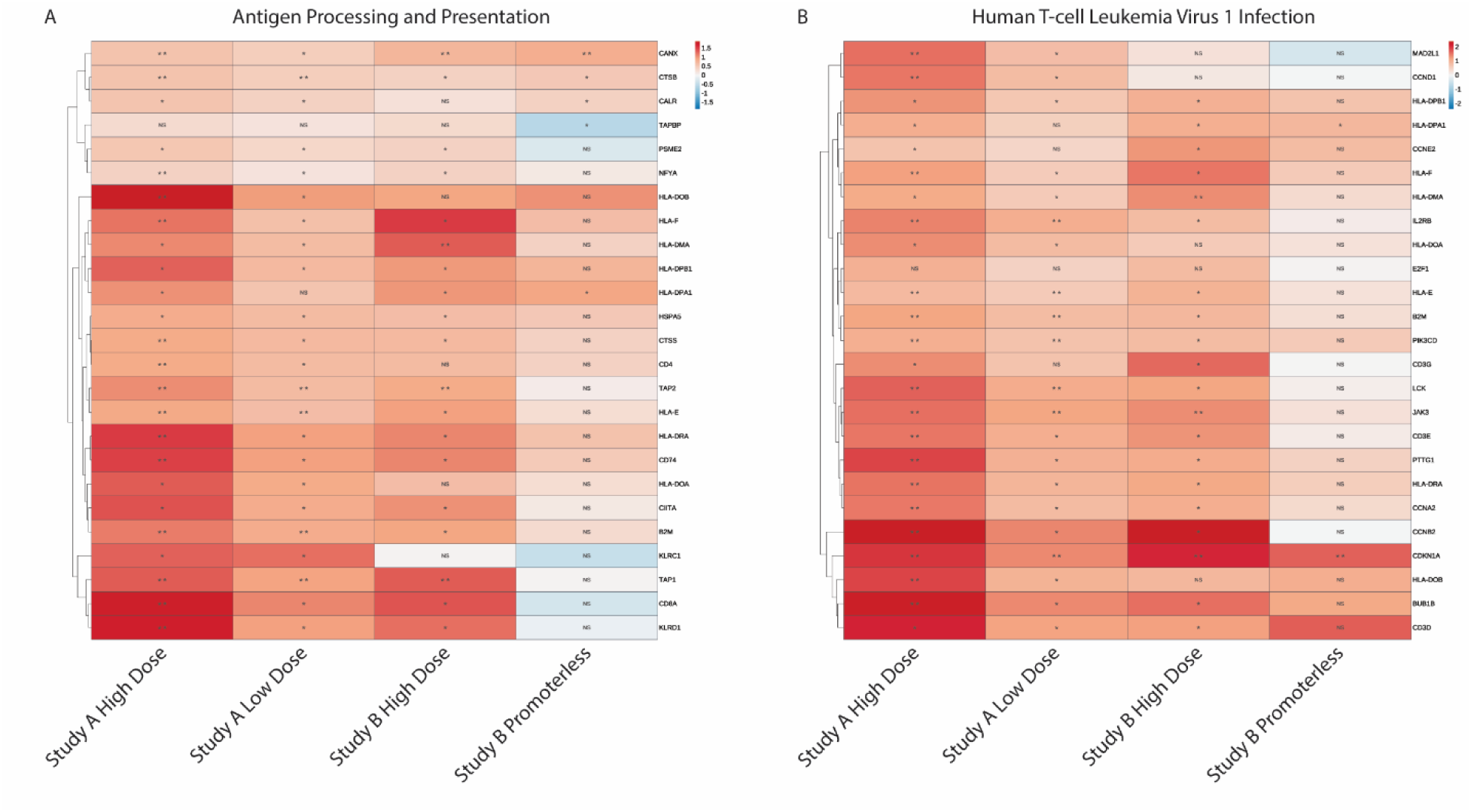
Significant pathways identified from differentially expressed genes in liver. The 25 most significant genes associated for each pathway are being displayed on the right of each table, segregated by study arm. The only two identified pathways up-regulated with the administration of scAAV9-CBA-GFP (fully functional vector) IT were: Antigen Processing and Presentation’ (A) and ‘Human T-cell Leukemia Virus 1 Infection’ (B). A few genes across both pathways were found to be also associated with the administration of the Promoterless vector. The heatmap colors represent log_2_FC and the dendrograms on the right depict hierarchical clustering. NA: Not Applicable; NS: Not Significant; *: p<0.05; **: p<0.01; ***: p<0.001

Taken together, we have observed a predominance of immunological genes associated with the interferon pathway for anti-viral responses in the DRGs as well as antigen presentation activity in the liver of non-human primates dosed with scAAV9 intrathecally. These results highlight a persistent low-grade innate immune response can be detected in this scAAV9-exposed tissue up to four weeks post-dosing. Such signature was absent in the groups exposed to empty capsid or the Promoterless vector, suggesting the expression of the transgene is key in that context.

### Individual gene signatures confirm that the most upregulated transcripts in organs at risk of toxicity predominantly play a role in immunological functions

To see the most prominent individual genes upregulated after intrathecal delivery of scAAV9 in the DRGs, we filtered for differentially expressed genes that were significantly dysregulated across studies A and B (scAAV9 full vector) compared to their respective vehicle control group, before sorting by largest log_2_FC values.

In the DRG, the most highly induced genes upon scAAV9 administration were immunoglobulin or interferon related genes (Figure 7A). Several genes in the top 25 list were seen in the pathway analysis results such as *GBP2*, and *CXCL10*. Others were observed in this top list such as *SLAM8* (cell surface protein of the CD2 family, involved in lymphocyte activation) and *PLAAT4* (phospholipase A1/2 and Acyltransferase 4), without a clear association with a broader biological pathway. Genes upregulated in the AAV group as well as weakly induced in the Promoterless group were also of interest, including *CXCR3* and *ZBP1*. Many of these significant genes are found to be upregulated through the interferon pathway.

**Figure 7:**
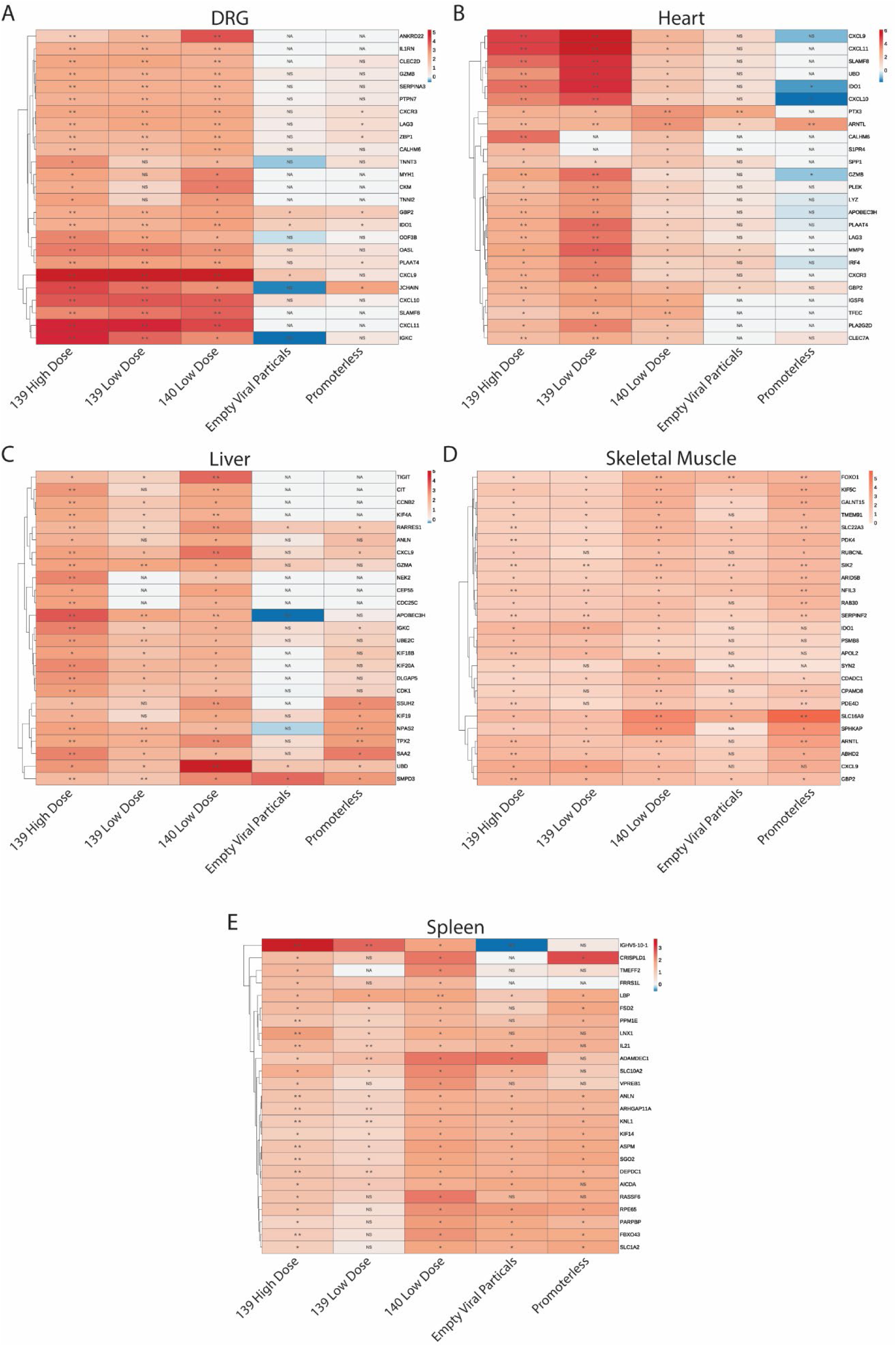
Highly Expressed Immunological Genes in AAV Dosed Groups. Heatmaps of the top 50 upregulated genes in high dose groups. Genes were grouped by hierarchical clustering. Color bar indicates log_2_FC. Characters within cells indicate significance. NA: Not applicable, NS: Not significant, *: P value < 0.05, **: P value < 0.01, ***: P value < 0.001.

In liver, 15 of the 25 genes were found to be uniquely significantly upregulated in scAAV9 dosed groups only (Figure 7C) as compared with the vehicle controls. Similarly, as our analysis done on the DRGs, many of these genes were immune related and included, for example, *TIGIT*, *GZMA*, and *IGKC*. Some familiar genes seen in the pathway analysis such as *CXCL9* were also seen in this top 25 list. Of interest, however, is the strong expression of *APOBEC3H* that was specifically detected in hepatic tissue and not in DRGs. *APOBEC3H* is a cytidine deaminase that plays a role in antiretroviral activity. Many genes identified were also related to the cell cycle such as *CCNB1*, *KIF18B*, and *CDC25C*. This may potentially indicate some form of liver regeneration following scAAV9 transduction (oval cell hyperplasia was one of the microscopic findings called for by the pathologist in those studies), which was not observed in the non-functional AAV dosed groups. It is important to note that some of these genes in the top 25, such as *CXCL9* and *IGKC*, are upregulated in the Promoterless group. However, many of these signals are more strongly upregulated in the groups given the full AAV viral particles expressing a transgene (GFP).

The heart, skeletal muscle and spleen, while not sites of primary toxicity, did share several upregulated genes with the DRG and liver in the top 25. Of note are genes associated with the interferon pathway such as *CXCL9*, and *IDO1* in the heart and skeletal muscles. Among them, *CXCL10*, *APOBEC3H* and *IRF4* were only observed in the heart (Figure 7B, D, E). Similarly, as DRG and liver, we have observed a majority of significantly upregulated genes in the heart of NHPs that have received the fully functional AAV (those genes were not significant in animals that have been dosed with non-functional AAV).

As we observed multiple similar genes upregulated in different organs, we have investigated what significantly upregulated genes were shared across the three organs of interest (Figure 8).

**Figure 8:**
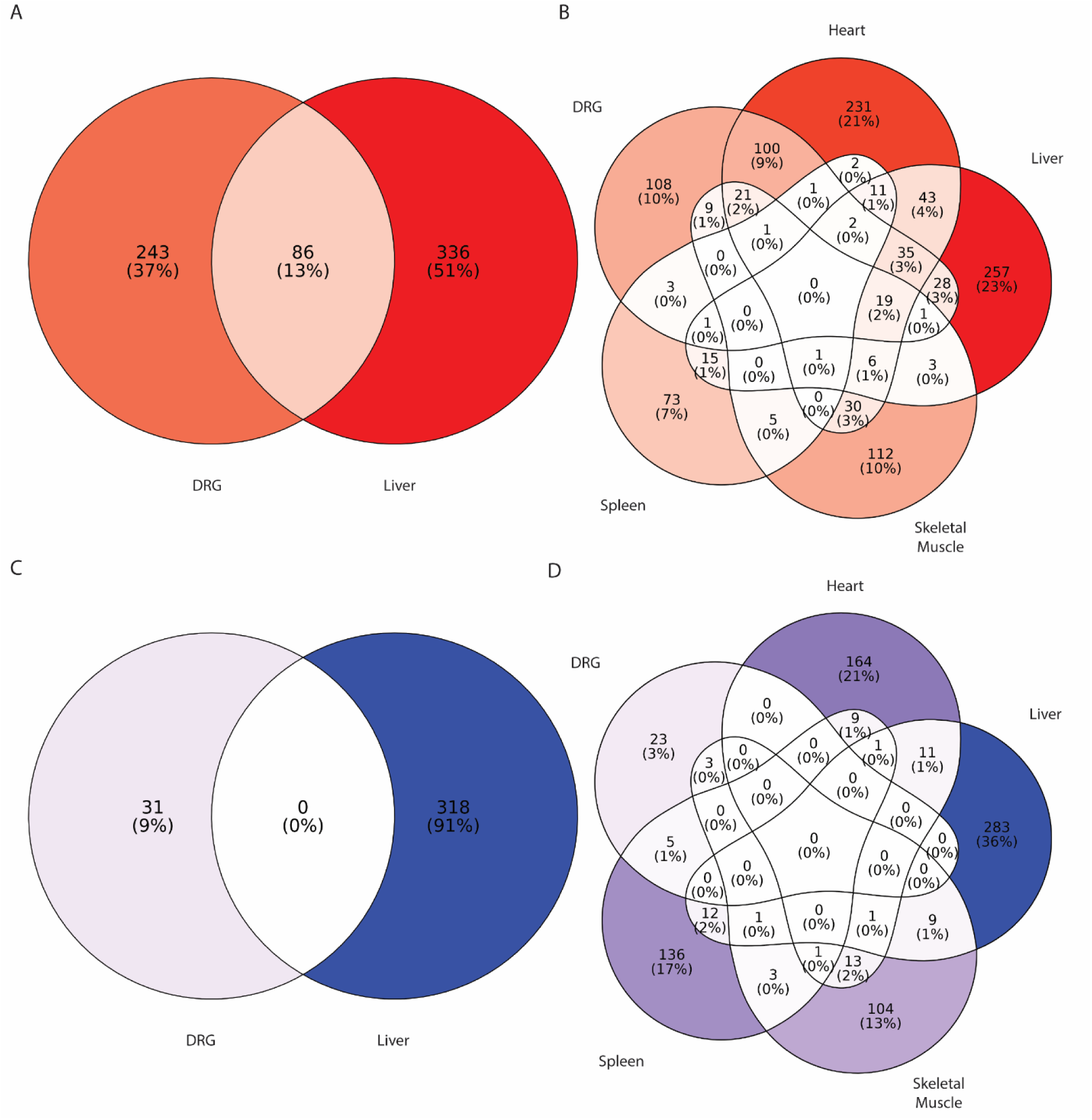
Shared Significantly Differentially Expressed Genes Common in Tissues with High Levels of Vector Genome. Venn diagrams of significantly upregulated genes shared between A) two or B) five tissues, and significantly downregulated genes across C) two or D) five tissues. Color intensity represents the number of significant genes. Number in loops indicate the number of genes. Percentages indicate the percentage out of all significant genes.

### Fewer AAV9-dependent downregulated genes are detected after intrathecal dosing with scAAV9 in non-human primates as compared to upregulated genes

Listing the top 25 downregulated genes in liver and DRGs revealed that fewer genes were distinctly repressed in the AAV dosed groups across studies A and B.

In the DRG, downregulated genes following administration of the fully functional AAV were related to neuronal development/organization (*NPAS1*), secretion (*LF*), or cell cycle progression (*S100A8* and *S110A9*). To the best of our knowledge, those genes are independent of one another and do not interplay in similar biological processes or pathways (Figure S3A). In the liver, only two genes*, NECAB2* (Receptor for 2A and metabotropic glutamate receptor modulator) and *SLC13A2* (citrate transporter), were found to be downregulated in the AAV dosed group. Similarly, only few genes were found to be significantly down regulated in the heart, skeletal muscle and spleen after intrathecal administration of scAAV9 (Figure S3B, D and E).

### Common immunological genes are upregulated in tissues where AAV-driven toxicity is reported

Investigating further the shared genes between the DRG and liver revealed it to be dominated by genes regulating immunological functions as well as the detection and response towards viral infections. While it is useful to analyze data through shared genes above a certain threshold, it is important to note that applying stringent cutoffs can result in biases to interpretations. For example, the expression of *CXCR3* is high in DRG and liver but its hepatic expression is slightly lower than the threshold (1.4 log_2_FC) and therefore *CXCR3* is not identified as a shared gene across both tissues. Abiding by this threshold, we have found many genes that were found in the top hits to be expressed across the organs (Table S7). Shared gene signatures such as *OASL*, *APOBEC*, *CXCL9*, and *IRF4* indicate that the interferon pathway is one pathway upregulated in the organs with the highest transgene expression. Similarly, T cell related genes such as *CCL5*, *CD3*, *CD8*, as well as B cell related immunoglobulin genes show an adaptive immune response is also being mounted in the organs of interest.

Additional comparisons between DRG & heart, DRG & liver, and heart & liver present with similarities, with many genes associated with interferon signaling (*CXCl10, 11,3, INFR1, 8*), TNF signaling (*TNFSF8, 13B*), B/T cell function (*ITK, CD1c, CD2, CD27*), and antigen presentation (*CD86, HLA genes, CSF2RA*). Showing similar immunological responses in various organs. Most notable was the clear lack of overlap in genetic signatures in the spleen and skeletal muscles compared to the other three tissues (Figure 8B). Further reinforcing the fact that there is commonality in responses in the tissues with the highest transgene expression.

Applying a similar approach to downregulated genes with a fold change lower than −1.4, the lack of overlap between the five tissue sites is striking (Figure 8D). Reducing the comparison to DRG and liver to get clearer picture was no more revealing, with no downregulated genes shared between the two tissue sources (Figure 8C).

## DISCUSSION

AAV-based gene delivery has led to the emergence of breakthrough therapies for the treatment of otherwise incurable genetic diseases. However, despite its efficacy, the risk of organ toxicity remains a limiting factor for its broader application to non-life-threatening disorders. To get a deeper insight into such toxicities, the work provides a detailed view of the sustained transcriptional changes following a single intrathecal administration of AAV9 in non-human primates. To our knowledge, this is the first time that an unbiased RNA sequencing analysis has been conducted across key AAV9-transduced tissues from cynomolgus macaques.

AAV9’s biodistribution in peripheral tissues after intrathecal administration in old-world macaques has been well characterized by us and other contributors in this field^12,29^, with the highest levels of vector genomes generally detected in the liver and DRG. Incidentally, AAV9-associated organ toxicities have been also mostly reported in those organs, as illustrated by the two present studies included in our analysis^10^. Interestingly, we also observed a limited correlation between the viral genome biodistribution and the transgene expression level detected across tissues, with the highest levels of transgene transcript detected in skeletal muscle and heart (the lowest levels of expression were found in spleen). This disparity between the load of viral genomes and the transgene product across tissues suggests that the fate of the viral genetic payload may vary across transduced cell types. As recently published by Hordeaux et al. in the context of liver transduction, such phenomenon appears to be dynamically regulated over the first few weeks post-dosing, leading to a drastic reduction in the level of transgene expression in hepatocytes with more limited impact on the viral genome^12^. Whether or not similar mechanisms are at play in DRGs or in other tissues (enabling a different output with regards to transgene product) is not known. Nonetheless, the fact that AAV9-associated organ toxicities happen in tissues with the highest level of viral genomes (but not transgene transcript) suggests that the amount of vector DNA entering the cells may be damaging and a key trigger of cellular stress rather than the transgene product itself, even if the expression of the transgene is also necessary to the observed changes (which are greatly reduced or even absent after dosing with empty capsids or the ‘Promoterless’ vector). The present work also highlights the long-term transcriptional impact of the viral vector following a single administration, which significantly affects many genes four weeks after dosing.

In the DRG, our analysis revealed a very distinct skewing of upregulated genes and relatively fewer downregulated genes after scAAV9-CBA-GFP IT administration. This was less evident in the liver, with a more balanced number of up- and down-regulated genes. In both tissues (DRG and liver), many of the most significantly induced genes were related to immunological functions, across both innate and adaptive immune systems (specifically, antigen presentation and interferon signaling). As highlighted throughout this manuscript, the number of transcriptionally dysregulated genes was much lower in animals dosed with empty capsids or the Promoterless vector, even though these experimental groups were still distinguishable from the vehicle control group.

Key immunological genes found to be upregulated following dosing with scAAV9-CBA-GFP belonged to the anti-viral interferon pathway, monocyte/macrophage functions, and T cell/B cell activity. The strong levels of key adaptive immune genes such as *CD3*, *CD40*, *CD80*, *CD86*, *TNFRSF13* (*APRIL*) and *TNFRSF13B* (*BAFF*) indicated not only the infiltration of T cells and B cells but also heightened activity of these cells. However, it is important to note that unlike patients that receive glucocorticoids, which are potent NF-κB inhibitors, no immunosuppressive regimen was provided to animals included in studies A or B. It is conceivable that such immune responses may be somewhat blunted in patients. The strong induction of genes belonging to the interferon signaling such as CXCL9, CXCL10, CXCR3, IRF proteins, and OAS suggests that this pathway may play an important role in AAV-driven organ toxicity. Previously published data have shown that CXCL9, CXCL11, and their receptor CXCR3, play a key role in the recruitment of T, NK and NKT cells into the liver and could contribute to hepatotoxicity^30^. Ajuebor M. and colleagues have reported that an infection by adenovirus could lead to an increase IFN-gamma levels in the liver, resulting in the up-regulation ^31^. In this study, researchers observed an influx of CXCR3^+31^. The role of CXCR3 in acute liver injury remains, however, relatively more controversial than other conventional pro-inflammatory chemokines. Indeed, Zaldicar et al. reported that *CXCR3*^30^. Interestingly, the researchers only observed this effect in *CXCR3* KO mice and not in *CCR1* or *CCR5* KO models. Whether protective or detrimental, evidence highlights a potential role of CXCR3 in liver toxicity. Another interesting observation from those results is the identification of type I and type II interferon signaling pathways, which mainly utilize the JAK/STAT pathway rather than NF-κB^32^. This could provide some explanation as to why glucocorticoids are not always as potent as expected in preventing AAV-driven toxicity. From those data we can hypothesize that interferon signaling is exacerbating the stress from AAV dosing on host cells to cause them to tip over to an inflammatory state and cause cellular toxicity.

Another key signature observed across liver and DRG is the induction of *CCR2* and *CCL19*, also known as monocyte chemotactic protein 1 receptor and macrophage inflammatory protein-3 beta respectively. These changes were accompanied by an increased expression of several genes tied to the Major Histocompatibility Complex Class II. Most prominent genes identified were CD74, HAL-DMB, in addition to components for the antigen loading such as TAP1 and TAP2. Together this may indicate a push for recruitment of macrophages in tissues exposed to high viral load.

In the cardiac tissue, the transcriptional changes observed were generally smaller in amplitude than what was observed in DRG and liver. Nonetheless, immune-related genes were also found to be up-regulated in this tissue even though no consistent pathological findings were observed. AAV-driven cardiotoxicity has recently been reported in the clinic ^24^, with two patients from Pfizer’s clinical trial (PF-06939926) and one patient partaking in Sarepta’s trial (SRP-9001) for Duchenne muscular dystrophy (DMD) presenting with myocarditis symptoms^25^. Another study attempting to treat DMD with “dead” *Staphylococcus aureus* Cas9 (d*Sa*Cas9) caused acute respiratory distress syndrome and cardiac arrest^15^. In those cases, it is conceivable that the disease context itself plays a significant role in the observed clinical signs and may not be recapitulated in healthy animals.

In summary, our work revealed a low-grade but long-lasting immunological response in known organs primarily targeted by AAV9. Specifically, responses observed in fully functional AAV groups that are tailored towards monocyte/macrophage, T, and B cells activity, and Type-II interferon responses. These results expand our understanding of the impact of such therapeutic vectors on tissues at peak of transgene expression and could provide novel avenues for mitigating AAV-driven organ toxicity.

## MATERIALS AND METHODS

### Animals

This analysis aggregates data from a total of 40 female cynomolgus macaques (Macaca fascicularis). Across two non-GLP toxicity studies (Table S1), 28 NHPs were dosed intrathecally with AAV9 viral vectors (including four NHPs administered empty capsid and four NHPs administered AAV9 viral vector with a Promoterless transgene) and 12 animals were administered with vehicle (tangential flow filtration [TFF] buffer). The animals were between 12 to 50 months old at dosing. All procedures were performed in compliance with the Animal Welfare Act, the Guide for the Care and Use of Laboratory Animals, and the Office of Laboratory Animal Welfare. Studies were approved by the Institutional Animal Care and Use Committee (IACUC) at the testing facilities where the studies were conducted. The primates were Asian-origin and initially outsourced from Envigo Global Services, Inc. (Alice, TX, USA). The in-life portion of those studies was performed at Labcorp (Madison, WI, USA). Animals were group-housed in European guideline (ETS 123)–compliant pens (up to three animals/pen), enriched with a bedding material when possible. Animals were given various cage-enrichment devices, including fruit, vegetables, or dietary enrichments. Prior to study assignment, animals were preselected based on their total anti-AAV9 antibody titers. Animals with the lowest titers were included in the study.

### AAV vectors: payload and manufacturing

Manufacturing and characteristics of the viral vectors used in the studies were detailed in previous publications^10,11,27^. In short summary, serotype 9 AAV viral vectors were used for both studies A and B. The genetic payloads of the vectors consisted of a self-complementary genome and the expression of the enhanced green fluorescent protein (eGFP) reporter transgene was driven by the cytomegalovirus (CMV) early enhancer and a hybrid CMV enhancer/chicken β-actin (CBA) promoter. For study B, four animals were also dosed with either a Promoterless payload packaged in AAV9 or empty capsids.

### General study design

Study A consisted of 24 female NHPs with three treatment arms: vehicle, scAAV-CB-GFP low dose (1.0×10^13^ vg/NHP), and scAAV-CB-GFP high dose (3.0×10^13^ vg/NHP). In study B, 16 females were divided into 4 treatment arms: vehicle, empty viral capsids, a ‘Promoterless’ scAAV9-CB-GFP with a scrambled CB promoter, and a fully functional scAAV9-CB-GFP (Table S1). All 4 arms received a dose of 2.73×10^13^ viral capsids/NHP. Both studies were conducted over 28 days and the test article was delivered intrathecally or intracisternal magna. In-life procedures during the two studies are detailed in our previous work^10^. Blood collection was conducted prior to AAV administration and/or on the day of administration and throughout the duration of the study ^10^. Intrathecal dosing procedures were also detailed in our previous report^10^. Briefly, the animal was anesthetized and animal had up to 1 mL of cerebral spinal fluid was withdrawn prior to a dose of 1.0×10^13^ vg/NHP or 3.0×10^13^ vg/NHP for study A, or 2.73×10^13^ viral capsids/NHP for study B. Dose volumes did not exceed 2 mL. At end of study, select tissue samples were collected and snap-frozen in liquid nitrogen for vector genome biodistribution analysis, RNA extraction and sequencing. Biodistribution of the vector genome was evaluated by digital droplet (dd) PCR on DNA isolated from collected tissue samples. scAAV9-CB-GFP vector genome were quantified with a C1000 Thermal Cycler (Bio-Rad, Hercules, California). For normalization, a two-copy reference gene (*CFTR*) primers and probe were used. Further details are published in Meseck *et al*^27^. Additional endpoints included histopathology and molecular localization across key transduced tissues, which was previously published^10^.

### Total RNA extraction from frozen tissue samples

Liver, heart, muscle, spleen, cervical DRG, thoracic DRG, and lumbar DRG tissues were selected for RNA sequencing. For cervical DRG, thoracic DRG, and lumbar DRG, RNA extraction was performed using the Qiagen RNeasy Plus Kit, accordingly to the manufacturer’s instructions. RNA extraction from liver, heart, muscle, and spleen tissues was done using the Qiagen RNeasy Universal Tissue Kit. After extraction, the RNA Integrity Number (RIN) was determined using the TapeStation 4200 by Agilent (RNA ScreenTapes and High Sensitivity RNA ScreenTapes) to evaluate the quality of the RNA extracts. In preparation for bulk RNA sequencing, extracted RNA samples were normalized using concentrations determined during QC and cDNA library construction was performed according to the Illumina Stranded mRNA Prep, Ligation (96 Samples) Kit. After cDNA library construction, DNA quality is checked using the TapeStation 4200 by Agilent (D1000 ScreenTapes). After normalization and pooling, RNA sequencing was performed using a NovaSeq6000 to obtain at least 20 million reads per sample. Collected data was analyzed according to the PISCES analysis package in DIEP and also in parallel raw counts data was collected in DIEP, exported to R Studio, and analyzed using the edgeR pipeline.

### Computational and statistical analysis

Bulk RNAseq Fastq files were mapped to the M. fascicularis ensemble gene file (V.5.0) using the Pisces Python package (V.0.1.5.1, Salmon V.1.3.0). Quantification of GFP transgene expression was conducted using a custom reference file based off the mfas V5.0 reference index file that included the GFP sequence. Results from mapped genes were analyzed using EdgeR (V.4.0.1) with R (V.4.3.2) using the following packages: ggVennDiagram_1.2.3, scales_1.2.1, BRGenomics_1.14.1, rtracklayer_1.62.0, GenomicRanges_1.54.1, GenomeInfoDb_1.38.1, janitor_2.2.0, readxl_1.4.3, ggbreak_0.1.2, shadowtext_0.1.2, ggpubr_0.6.0, reshape2_1.4.4, showtext_0.9-7, showtextdb_3.0, sysfonts_0.8.9, org.Hs.eg.db_3.18.0, AnnotationDbi_1.64.1 IRanges_2.36.0, S4Vectors_0.40.1, Biobase_2.62.0, BiocGenerics_0.48.1, pheatmap_1.0.12, RColorBrewer_1.1-3, ggrepel_0.9.4, magrittr_2.0.3, edgeR_4.0.1, limma_3.58.1, lubridate_1.9.3, forcats_1.0.0, stringr_1.5.1, dplyr_1.1.3, purrr_1.0.2, readr_2.1.4, tidyr_1.3.0, tibble_3.2.1, ggplot2_3.4.4, tidyverse_2.0.0, clusterProfiler_4.10.1, BiocManager_1.30.22, biomaRt_2.58.0. All analysis in EdgeR including pathway analysis were run under recommended settings unless otherwise noted.

## Supporting information

Supp_Fig_1_Biodistribution_and_Transgene

Supp_Fig_2_SkelMusc_Volcano

Supp_Fig_3_Spleen_Volcano

Supp_Fig_4_Top_25_Down

SuppTable_1_Study_Design

SuppTable_2_DRG_MFas_Merged_KEGG_Table

SuppTable_4_Liver_MFas_Merged_KEGG_Table.csv - Shortcut

SuppTable_3_Heart_MFas_Merged_KEGG_Table

SuppTable_5_Muscle_MFas_Merged_KEGG_Table

SuppTable_6_Spleen_MFas_Merged_KEGG_Table

## Data Availability Statement

All analyses included in this manuscript are available (and its supplemental information files). Raw sequencing data can be shared upon request.

## Acknowledgments

We thank Cameron McElroy and Jayson Chen for their support as study monitors for the NHP studies conducted.

## Author Contributions

**FA** led the analyses of the RNAseq data and primarily contributed to writing the manuscript. **EH** contributed to the study design, sample preparation, data analysis and interpretation as well as writing the manuscript. **MM** performed the sample preparation (RNA extraction), QC check and library preparation for RNA sequencing. **VR, KB** and **JS** contributed to the sequencing and data QC. **TDR** performed biodistribution for studies presented in this work. **MOS** and **PC** provided support for the development of the analytical pipeline and gave input on data analyses. **FA, MM**, **VR, KB, JS, MOS, PC**, **KM**, and **EH** all reviewed the manuscript.

## Declaration of Interests Statement

**All co-authors** are employees of Novartis Institutes for BioMedical Research and several authors own Novartis stock or other equities.

## Notes

### Competing Interest Statement

All co-authors are employees of Novartis Institutes for BioMedical Research or have been employed by Novartis and several authors own Novartis stock or other equities.

### Funding

- Novartis (United States)

