## Supplementary figures and images for "Transcriptional changes in non-human primate tissues after intrathecal delivery of serotype 9 adeno-associated viral vector: insights into organ toxicities"

### Supp_Fig_1_Biodistribution_and_Transgene

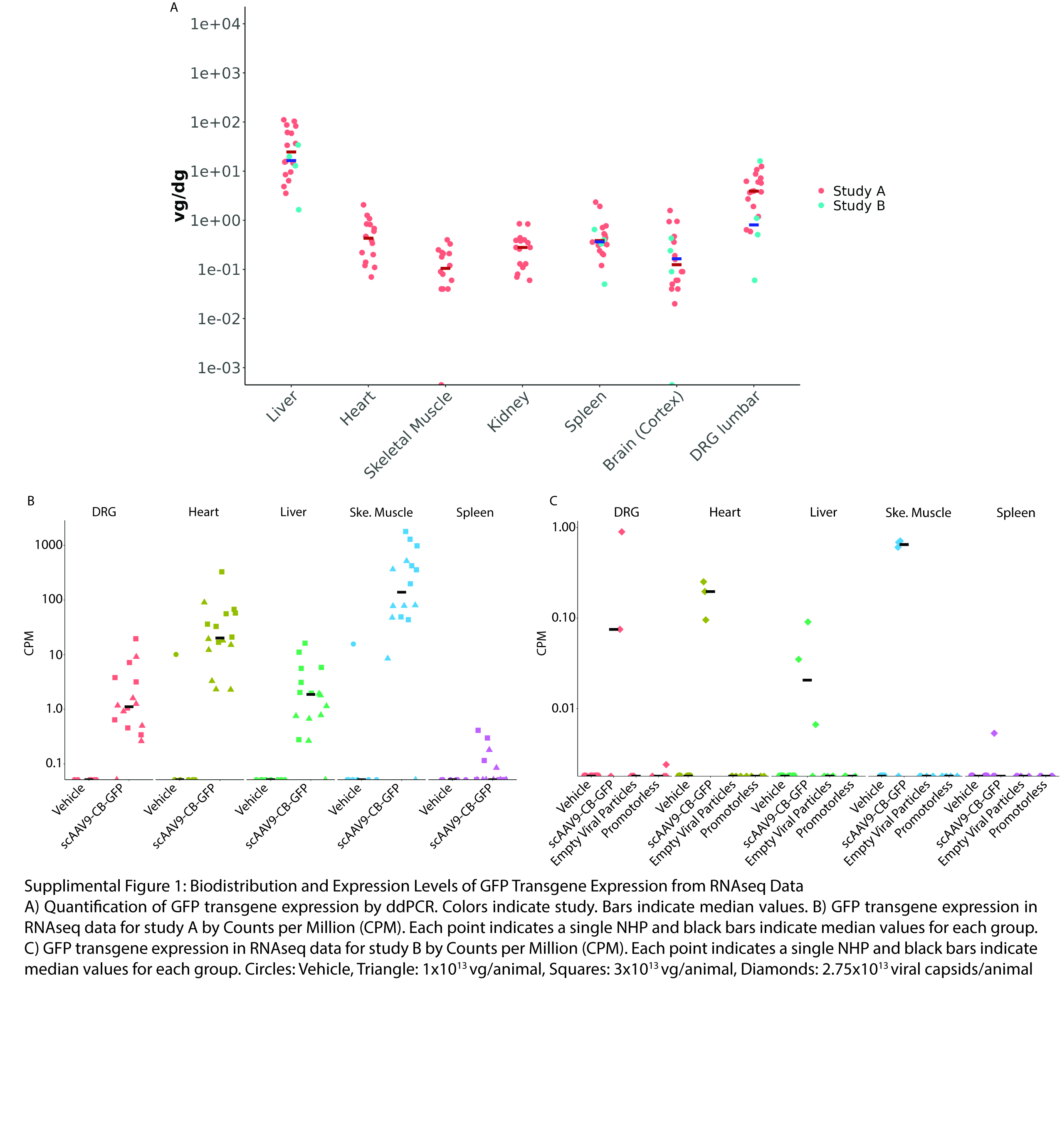

### Supp_Fig_2_SkelMusc_Volcano

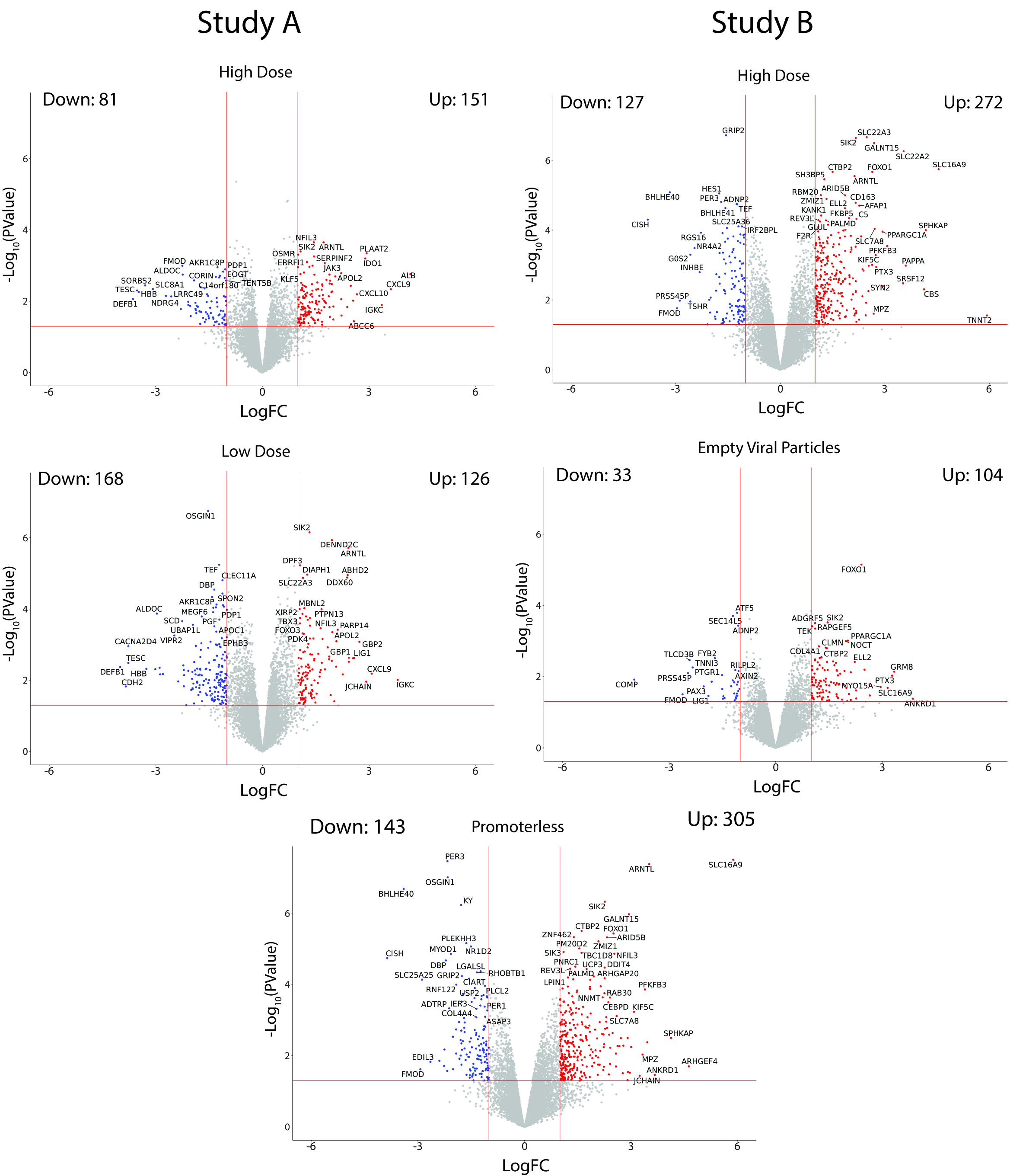

### Supp_Fig_3_Spleen_Volcano

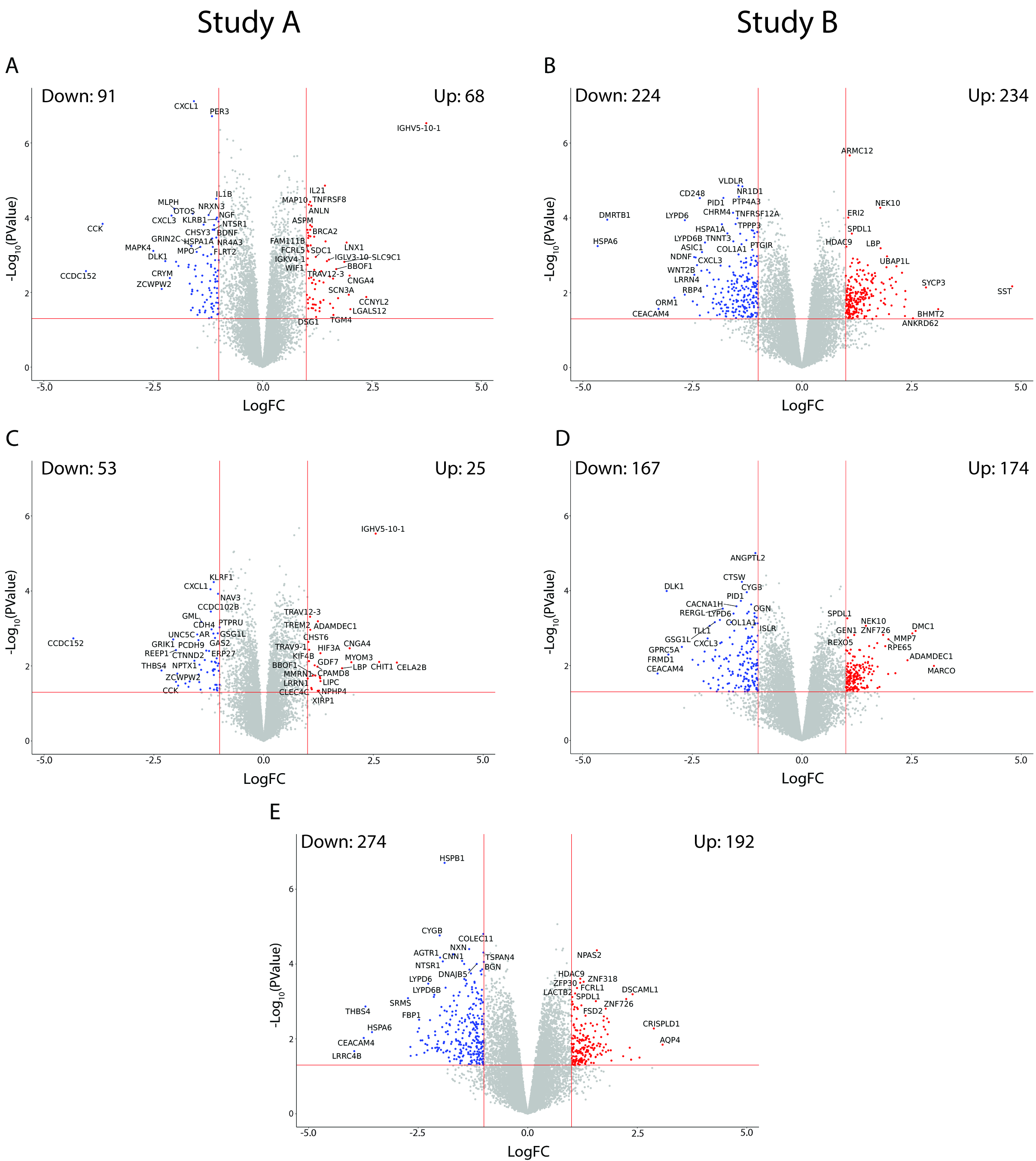

### Supp_Fig_4_Top_25_Down

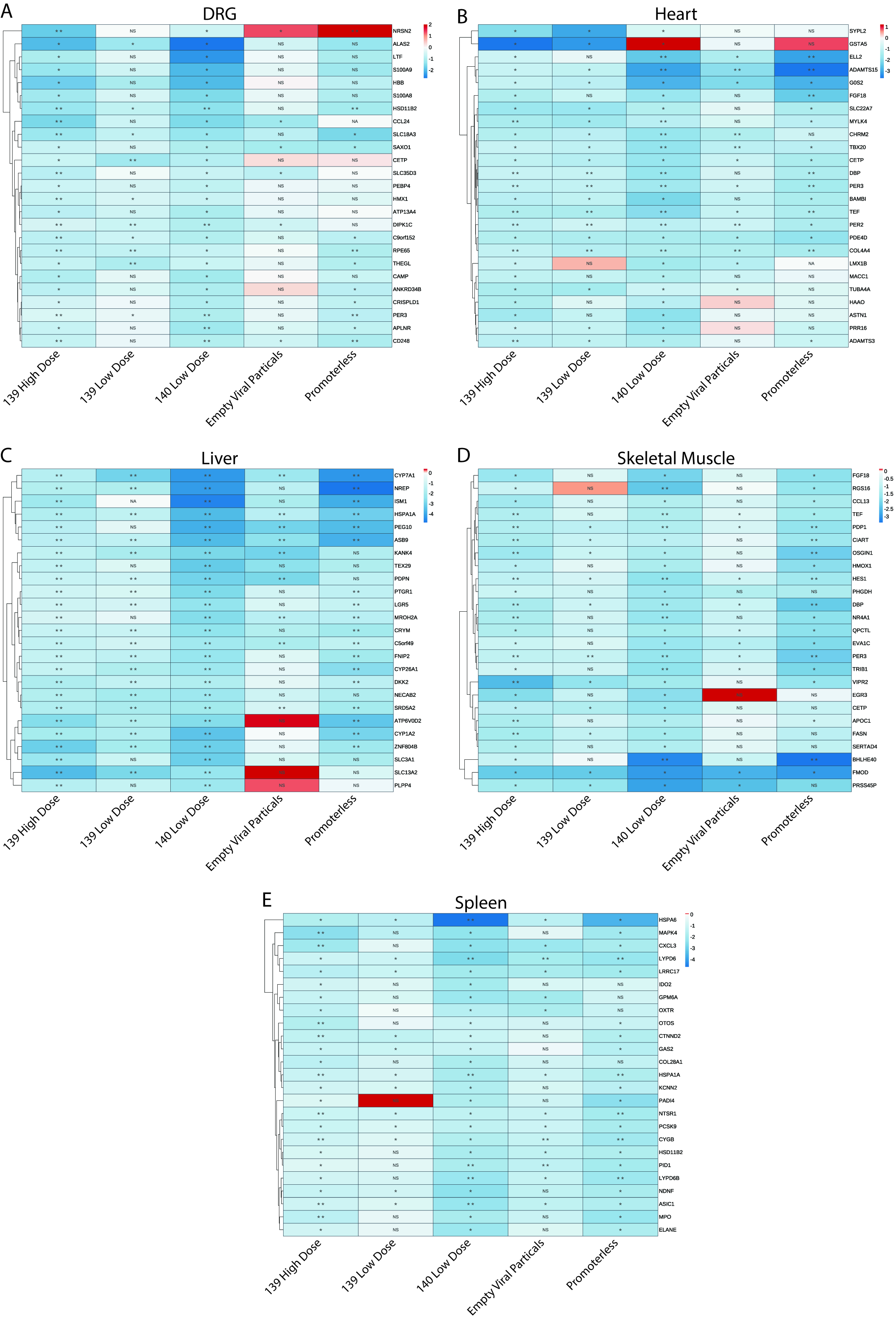
